# THE TRANSCRIPTIONAL RESPONSE TO OXIDATIVE STRESS IS INDEPENDENT OF STRESS-GRANULE FORMATION

**DOI:** 10.1101/2021.08.18.456454

**Authors:** Amanjot Singh, Arvind Reddy Kandi, Deepa Jayaprakashappa, Guillaume Thuery, Devam J Purohit, Joern Huelsmeier, Rashi Singh, Sai Shruti Pothapragada, Mani Ramaswami, Baskar Bakthavachalu

**Affiliations:** National Centre for Biological Sciences, TIFR, Bangalore 560065, India; Tata Institute for Genetics and Society Centre at inStem, Bellary Road, Bangalore 560065, India; Trinity College Institute of Neuroscience, School of Genetics and Microbiology, Smurfit Institute of Genetics and School of Natural Sciences, Trinity College Dublin, Dublin-2 Ireland; School of Basic Sciences, Indian Institute of Technology, Mandi 175005, India

**Keywords:** Oxidative stress response, stress granule, RNA-binding protein, transcriptome, chaperone, HSP, G3BP, Rasputin, Ataxin-2, *Drosophila*, S2 cells

## Abstract

Cells respond to stress with translational arrest, robust transcriptional changes, and transcription-independent formation of mRNP assemblies termed stress granules (SGs). Despite considerable interest in the role of SGs in oxidative, unfolded-protein, and viral stress responses, whether and how SGs contribute to stress-induced transcription has not been rigorously examined. To address this issue, we characterized transcriptional changes in *Drosophila* S2 cells induced by acute oxidative-stress and assessed how these were altered under conditions that disrupted SG assembly. Sodium-arsenite stress for 3 hours predominantly resulted in the induction or upregulation of stress-responsive mRNAs whose levels peaked during cell recovery after stress cessation. The stress-transcriptome is enriched in mRNAs coding for protein chaperones, including HSP70 and low molecular-weight heat shock proteins, glutathione transferases, and several non-coding RNAs. Oxidative stress also induced prominent cytoplasmic stress granules that disassembled 3-hours after stress cessation. As expected, RNAi-mediated knockdown of the conserved G3BP1/ Rasputin protein inhibited stress-granule assembly. However, this disruption had no significant effect on the stress-induced transcriptional response or stress-induced translational arrest. Thus, SG assembly and stress-induced effects on gene expression appear to be driven by distinctive signaling processes. We suggest that while SG assembly represents a fast, transient mechanism, the transcriptional response enables a slower, longer-lasting mechanism for adaptation to and recovery from cell stress.

## INTRODUCTION

Oxidative stress can have several cellular consequences, including DNA damage and increased levels of oxidized and misfolded proteins (Schieber and Chandel, 2014). It also activates components of the cellular integrated stress response (ISR) pathway, including stress kinases that modify the mRNA translational machinery (Balchin et al., 2016; Costa-Mattioli and Walter, 2020; Gidalevitz et al., 2011). Phosphorylation of the eukaryotic initiation factor, eIF2α, results in translational inhibition together with the formation of stress granules (SG), assemblies of translationally arrested mRNAs, RNA-binding proteins and accessory components (Kedersha et al., 1999; Kedersha and Anderson, 2002; Ron, 2002).

Pathways and proteins involved in the ISR have been implicated in normal aging and in neurodegenerative disease (Halliday et al., 2017; Krukowski et al., 2020; Radford et al., 2015). Increased levels of oxidative stress are also thought to be associated with normal brain aging (Milton and Sweeney, 2012). Consistent with this, unusual SG-related neuronal inclusions have been observed in post-mortem brain samples of aged but not young brains (Bäuerlein et al., 2017; Geser et al., 2010; Ginsberg et al., 1998). SGs have gained even more significance since the discovery that protein inclusions associated with neurodegenerative diseases can contain SG components. In some cases, both inclusion formation and disease progression depend on factors that drive normal SG assembly (Advani and Ivanov, 2020).

In addition to SG formation, oxidative stresses regulate transcription factors such as FOXO, HSF1, and Nrf2 to induce changes in the cellular transcriptome (Donovan and Marr, 2015; Doonan et al., 2019; Fedoroff, 2006; Vihervaara et al., 2018). In particular, stress increases the expression of mRNAs coding for cytoprotective proteins, including protein chaperones and modulators of lipid oxidation (Jacobson et al., 2012). The third effect of acute oxidative stress is to induce translational arrest for the majority of cellular mRNAs. Here we ask whether these different stress responses occur independently of each other. In particular, we test whether signaling mediated through assembled stress granules contributes to the transcriptional responses to stress as has been suggested by the role of SGs in signaling required for transcription of genes involved in viral defense (Fung et al., 2013; McCormick and Khaperskyy, 2017; Tsai and Lloyd, 2014). In cultured *Drosophila* S2 cells, we: (a) document the requirements and kinetics of SG formation and disassembly; (b) obtain robust data sets for stress-induced transcriptional changes during and after acute stress; and (c) examine how the stress-induced transcriptome and global mRNA translation is altered when SG assembly is perturbed. We address these issues in *Drosophila* cells, partly for the ease with which stress-granule assembly can be visualized and perturbed in these cells but mainly because *Drosophila* allows facile, future follow-up experiments to assess the function of stress-regulated genes *in vivo*.

As anticipated, *Drosophila* S2 cells acutely exposed to the well-known stressor sodium arsenite show robust formation of Ataxin-2 and Rasputin (Rin)/Ras GTPase-activating protein-binding protein 1 (G3BP1) positive SGs along with simultaneous inhibition of global translation (Escalante and Gasch, 2021; Ivanov et al., 2019; Jain et al., 2016; Kedersha et al., 2016; Kedersha and Anderson, 2007; Wheeler et al., 2016). Parallel RNA-seq analyses show that arsenite stress also induces upregulation of around 300 different transcripts. Following three hours of post-stress recovery in the absence of arsenite, SGs disassemble and become invisible. In contrast, the vast majority of stress-induced mRNAs remain upregulated, consistent with a model in which SGs represent an acute protective mechanism that provides cells time to launch a longer-lasting transcription-dependent program for recovery from stress. In cells lacking Rin, although stress granules are not visible, stress-induced translational arrest and stress-induced transcription remain unchanged. These data indicate stress-granule formation is largely dispensable for oxidative-stress-induced changes to gene regulation.

## RESULTS

### Kinetics of assembly of arsenite-induced stress granules in *Drosophila* S2 cells

To understand cellular changes occurring during oxidative stress and subsequent recovery, we employed sodium arsenite as a stressor in *Drosophila* S2 cells, an established cellular model for studying the stress response (Aguilera-Gomez et al., 2017; Farny et al., 2009). Consistent with prior observations (Bakthavachalu et al., 2018; Farny et al., 2009), we found that exposure of cells to 0.5mM arsenite for 1h leads to the formation of numerous Ataxin-2 (Atx2) and Rin/G3BP positive stress granules; these appeared larger and more distinct after 3h of stress (Fig. 1A-i,-ii). To determine the temporal dynamics of clearance of SGs, we stressed the cells for 3h, allowed them to recover by replacing the stressor with a fresh culture medium, and monitored SGs at specified time points afterward. Although some cells still had several granules, recovery from stress, in general, was accompanied by the progressive disappearance of SGs with majority of cells having no or a few granules (Fig. 1B; Supplementary Fig. 1A). While some Atx2-positive granules remained after 1h of recovery (Fig. 1A- iv), none were visible after 3 hours in most cells (Fig. 1A-v).

**Fig. 1.**
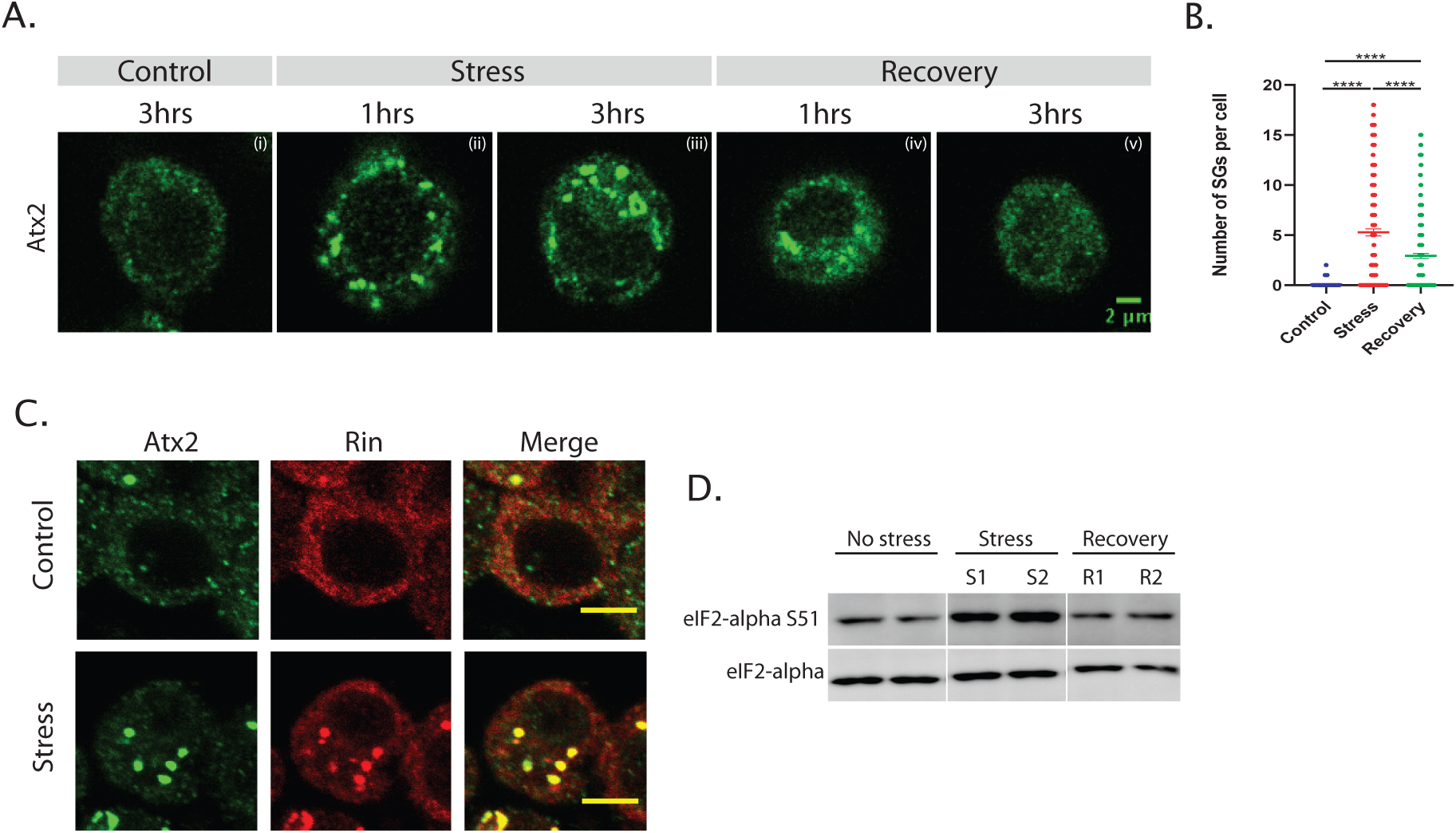
Kinetics of assembly of arsenite-induced SGs in *Drosophila* S2 cells. **A.** Progression of arsenite-induced SGs assembly. Untreated S2 cells do not show any granular structures stained by anti-Atx2 antibodies. Atx2-positive stress granules appear within 1-hour of arsenite exposure. More distinct granules are seen after 3 hours. Upon removing stress, the granules gradually start to clear, and after 3h of recovery, Atx2 returns to its normal diffused state. Staining was performed using antibodies against Atx2. **B.** Number of granules present per cell under control, stress and recovery are plotted. The number of cells and the granules present in the cells were quantified using CellProfiler. Mann-Whitney U-test shows that there was a significant difference in the number of granules between stressed and recovered cells (p < 0.05). Images and raw values corresponding to the analyses are shown in Supplementary Fig. 1A and Supplementary File 1. **C.** Atx2 and Rin co-localize in SGs, shown by staining with antibodies against Atx2 and Rin. **D.** Western blotting of total cell lysates shows that eIF2-α is hyper-phosphorylated during 1h (S1) and 3h (S2) stress. Cells were allowed to recover for 3h after both 1h (R1) and 3h (R2) of stress. Total eIF2 α levels do not show any change. Uncropped Western blots are shown in Supplementary Fig. 1B. Scale bar represents 2μm (A) and 10μm (C).

To address whether the disappearance of stress granules after recovery correlated with reduced stress signaling, we assessed phosphorylation levels of eIF2α at S51 (Fig. 1D). It is well established that stress-kinases such as PEK and GCN2 phosphorylate eIF2α trigger arsenite-induced SGs formation (Farny et al., 2009). Consistent with this, we observed that eIF2α phosphorylation in S2 cells is significantly elevated following either 1h or 3h of exposure to arsenite (Fig. 1D). After 3 hours of arsenite removal, levels of eIF2α phosphorylation were comparable to those under control conditions: moreover, there was no change in total eIF2α expression under any of these conditions (Fig. 1D). Taken together, these observations confirm and extend previous findings in S2 cells, showing that oxidative-stress induced SGs are transient, dynamic structures whose assembly/disassembly is concomitant to eIF2**α** phosphorylation and whose formation is associated with the shutdown of protein translation.

### Distinctive acute stress and post-stress (“recovery”) transcriptomes

Genes transcriptionally regulated by stress could potentially encode factors involved in regulating the assembly and clearance of SGs or managing molecular or physiological consequences of stress. To identify molecules potentially involved in these processes, we examined transcriptional changes in S2 cells under acute stress conditions and following recovery. We isolated total RNA from cells that were (a) untreated, (b) stressed for 3h, and (c) recovered for 3h following 3hr stress and used RNA-Seq to identify and analyse polyA-selected RNA populations in each condition (Fig. 2A). Three independent biological replicates were used for each of the three conditions. A total of more than 114 million high-quality reads (average ∼10 million reads per sample) were generated and mapped to the *Drosophila* genome using STAR v2.5.3 (Supplementary File 2). The uniquely mapped reads for each sample were processed using HTSeq to determine the transcripts’ normalized expression levels. The correlation coefficient values demonstrate high similarity (0.992 to 1.0) across the biological replicates and clear differences in global transcriptomes during normal, stress, and recovery conditions (Fig. 2B; Supplementary Fig. 2). Thus, the analyses show that control transcriptomes differ significantly from those of cells during stress and following 3 hours of recovery.

**Fig. 2.**
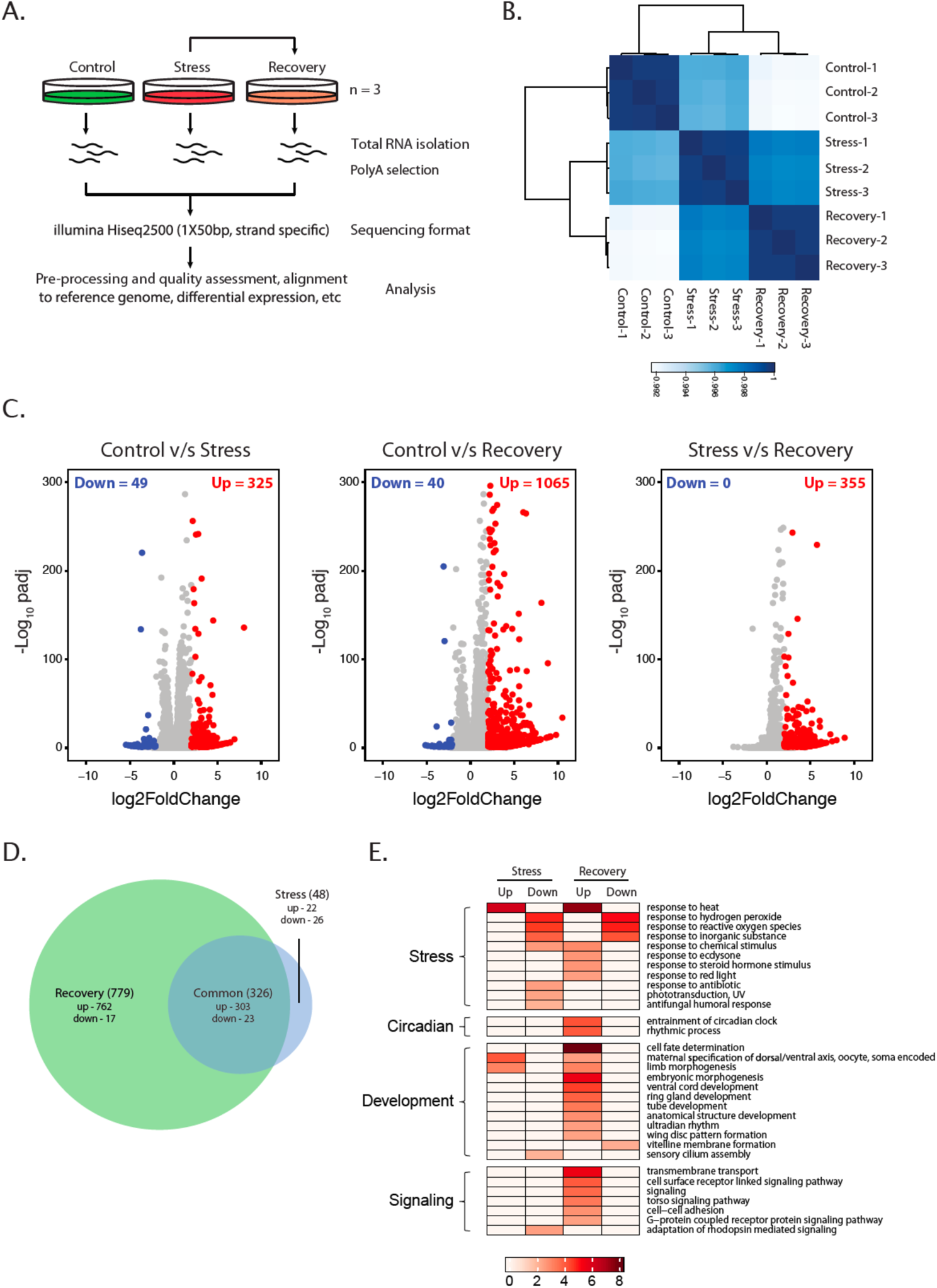
Distinctive normal, stress, and recovery transcriptomes. **A.** Schematic representation of the experimental design. Cells were stressed for 3h with 0.5mM sodium arsenite and pelleted for RNA isolation. For Recovery, arsenite was removed after 3h of stress, and cells were washed three times with S2 cell culture media and then maintained in fresh media for an additional 3h. Cells were subsequently harvested for RNA isolation. **B.** Pearson’s correlation plot visualizing the correlation between samples. The colour scale represents the range of correlation coefficients displayed. **C.** Volcano plots showing the differentially regulated transcripts in stress and recovery. **D.** Venn diagram depicting the overlap between “stress” and “recovery” transcriptomes. **E.** Gene ontology analysis of differentially expressed genes during stress and recovery. The enriched GO terms (biological process) in differentially expressed (up/down) genes under stress and recovery conditions compared to control conditions are shown via heatmap. The scale at the bottom represents enriched GO terms in –log10p-value.

### Strong transcriptional changes are observed after stress cessation

To identify the main differences in transcriptomes across cells at rest, under stress, and after recovery, we identified genes whose expression was altered at least log2 fold change of 2 with an adjusted P-value (padj) <0.05 between conditions (using the average expression values across replicates in each). Of 374 transcripts that were differentially regulated after 3 hours of stress, we found that levels of 325 transcripts were elevated and only 49 reduced compared to untreated cells (Fig. 2C, Supplementary File 3), indicating that stress predominantly resulted in induction of transcription.

Transcriptomes of cells 3-hours after recovery were even more different from untreated cells, than were transcriptomes of cells 3-hours after stress. Thus, 1105 transcripts showed at least log2 fold change of 2 difference in expression in cells 3-hours post-recovery compared to untreated cells. Of these 1105 transcripts, 1065 were upregulated, and 40 transcripts were downregulated (Fig. 2C). More detailed comparisons indicate that mRNAs upregulated more than log2 fold change of 2 after 3-hours recovery were generally induced, albeit to a lesser extent, after stress alone. Consistent with this, when transcriptomes of cells 3-hours post-recovery compared with transcriptomes of stressed cells, we found only 355 transcripts that showed a log2 fold change of 2 increase in expression after 3-hours of recovery (Fig. 2C). Intriguingly, mRNAs induced by acute stress stayed upregulated for hours after the stressor was removed. Thus, the expression of almost all the transcripts differentially regulated in stress was also similarly altered following 3 hours of post-stress recovery (Fig. 2D). Only 48 transcripts were unique to stress transcriptome; 22 of these were upregulated while 26 were downregulated. These observations clearly show that unlike SGs, which disassemble when the stressor is removed, stress-induced transcriptional changes persist long after the stressor is gone.

A Gene Ontology (GO) enrichment analysis provided a high-level view of functional classes of genes over-represented during stress and subsequent recovery (Fig. 2E). In particular, mRNAs known to respond to increased temperature and heat stress were particularly highly enriched during stress and after a 3-hour recovery (Fig. 2E, Supplementary File 3).

### Multiple classes of potentially cytoprotective mRNAs induced by stress

A detailed analysis of the identity of stress-regulated mRNAs was consistent with a model in which oxidative stress predominantly leads to the upregulation of a cohort of genes required for a delayed response to acute stress in S2 cells (Fig. 2C, D, Supplementary Fig. 3). Of 100 genes most strongly upregulated after 3 hours of recovery, several encoded heat-shock proteins (HSP) of the HSP70 (Hsp70Bc, Hsp70Bbb, Hsp70Ba, Hsp68), HSP40 (DnaJ-1), low molecular weight (LMW), HSP (Hsp23, Hsp26, Hsp27) families, and co-chaperones (stv) families (Fig. 3A). Interestingly, several of these upregulated genes have been previously shown to be regulated by heat stress in *Drosophila* (Vos et al., 2016). This indicates significantly overlapping cellular mechanisms for the management of oxidative stress and heat stress, which is consistent with previous observations for the phenomenon of “cross-tolerance” in other organisms (Mittal et al., 2012; Perez and Brown, 2014; Vert and Chory, 2011). The upregulation of *Hsp* mRNAs seems to be an evolutionarily conserved response required for folding the misfolded/aggregated proteins during stress (Verghese et al., 2012). In this regard, the upregulation of LMW HSPs (Hsp20/α-crystallin family) and HSP70 mRNAs upon oxidative stress suggests extensive misfolding of proteins under the conditions, which needs to be managed such that they can be refolded into the native state for the cell to recover. Small HSPs (sHSPs) function as “holdases” and prevent the formation of denatured protein aggregates in the cell and while HSP70s are the main folding agents of nascent polypeptide chains as well as for misfolded proteins during periods of stress (Finka et al., 2016; Vos et al., 2016). HSP27 has also been shown to bind poly-ubiquitin chains and interact with 19S proteasome (Bozaykut et al., 2014; Mogk et al., 2019), suggesting elevated levels of this protein during recovery may also play a role in protein triage.

**Fig. 3.**
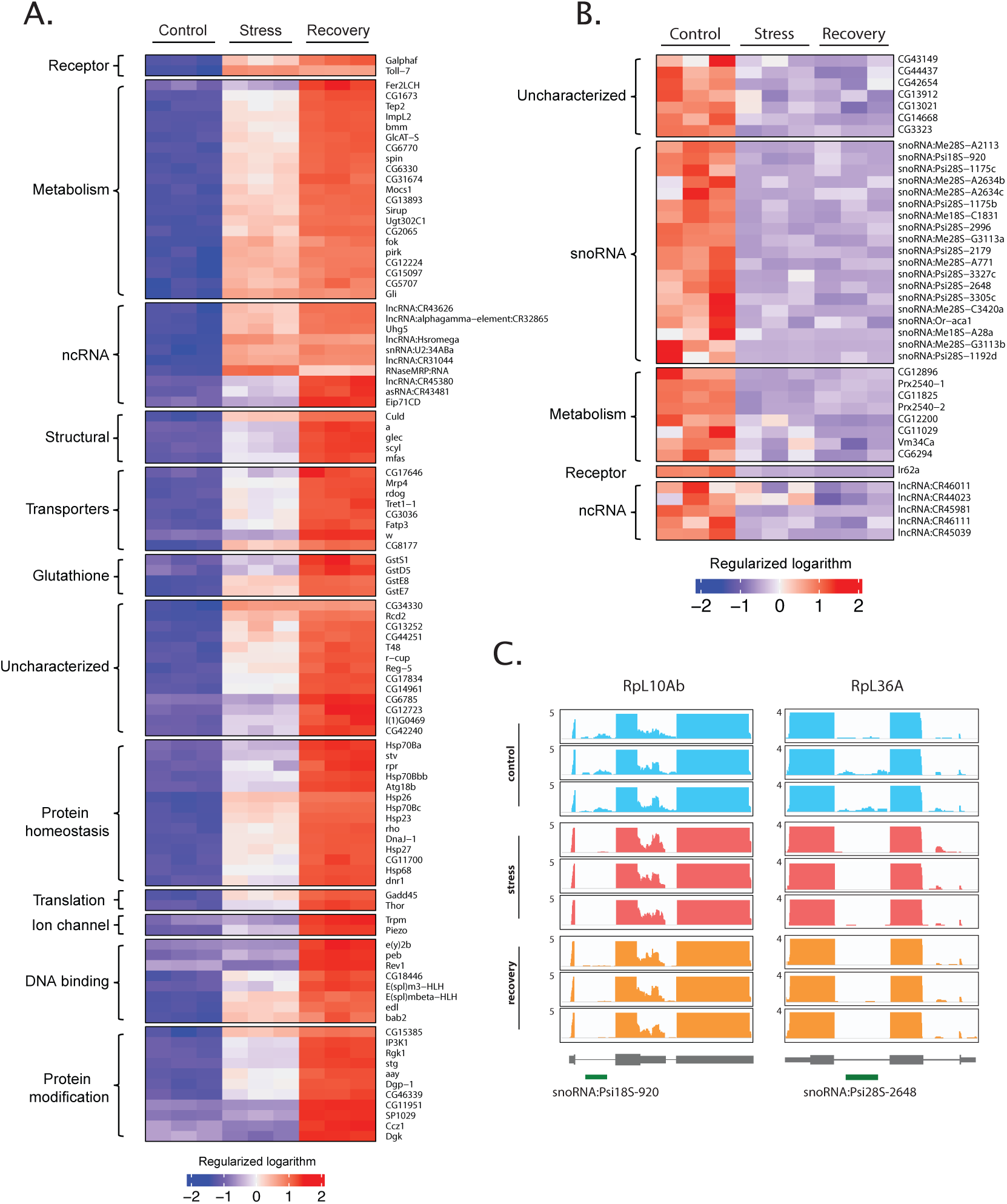
Oxidative stress results predominantly in the induction of target mRNAs. **A.** Heat map for 100 genes most robustly upregulated following 3-hours of recovery from 3-hours of acute stress. Genes are grouped based on predicted cellular functions. The fold induction is indicated in the colour scale below. **B.** Heat map shows a smaller group of mRNAs for which reads are substantially decreased after acute stress. The colour scale bar indicated fold changes represented. **C.** Bed graphs showing the reads under control, stress, and recovery corresponding to the parent genes RpL10Ab and RpL36A, which harbour snoRNA:Psi18S-920 and snoRNA:Psi28S-2648.

In addition, we noticed a more specific upregulation of transcripts encoding factors expected to help counter the effects of oxidative stress (Fig. 3A), in particular, genes for glutathione-S-transferases (GstD5, GstE7, GstE8, and GstS1). GSTs are detoxification enzymes that detoxify reactive oxygen species by catalyzing the addition of glutathione (GSH) and protect the cell from oxidative damage (Mailloux et al., 2013).

Apart from Hsp and GST transcripts, several non-coding (nc) RNAs, CR43481, CR45380, Uhg5, CR31044, CR43626, CR32865, Hsr-omega, and RNaseMRP:RNA were also upregulated (Fig. 3A). We speculate that these, as well as upregulated mRNAs encoding DNA-binding proteins like bab2, edl, e(y)2b, peb, Rev1, E(spl)m3-HLH, and E(spl)mbeta-HLH could potentially regulate the expression of “late” genes, such as those strongly induced 3-hours of recovery, of which several interestingly encode metabolic factors (Fig. 3A, Supplementary Fig. 3).

An unexpected finding is that reads corresponding to several small nucleolar RNA (snoRNA) genes that are frequent in resting cells are highly reduced in number both during stress and after 3 hours of recovery (Fig. 3B, Supplementary Fig. 3B). snoRNAs are RNA PolII-transcribed, short essential non-protein-coding RNAs (60-300 nucleotides long) that are mostly localized to nucleoli (Bratkovič et al., 2020; Kufel and Grzechnik, 2019). The primary function of snoRNA-ribonucleoprotein complex is post-transcriptional maturation of ribosomal RNA (rRNA) and small nuclear RNAs (snRNA) through 2-O’-methylation and pseudouridylation. Because most snoRNAs do not have polyA tails, it was quite surprising to find reads corresponding to snoRNAs in our polyA libraries under normal conditions. Since several snoRNAs are encoded in the introns of pre-mRNAs, particularly those encoding ribosomal proteins (Bratkovič et al., 2020; Kufel and Grzechnik, 2019), one possibility is that RNA-Seq reads for snoRNAs correspond to the introns from unspliced, polyadenylated nuclear pre-mRNAs (Supplementary Fig. 3C). We therefore, examined whether the reduced number of snoRNA reads after stress could correspond to increased splicing of the parent pre-RNAs.

The bed graph files for two intron-encoded snoRNAs; snoRNA:Psi18S-920 and snoRNA:Psi28S-s648 (Fig. 3C), show that RNA-Seq reads corresponding to these snoRNAs are almost absent during stress and recovery. If this decrease corresponds to reduced transcription, then the parent mRNAs must also be downregulated. Instead, the normalized counts of 11 such parent ribosomal protein genes show that in contrast to respective snoRNA reads, their levels are slightly elevated, certainly not decreased, during stress as well as after 3 hours of recovery (Supplementary Fig. 3C). Similarly, transcript levels of snoRNA:Me28S-A2113 and snoRNA:Psi28S-2996, which arise from RpL30 and RpL5 respectively, also show significant reduction both during stress and recovery (Supplementary Fig. 3D). The most likely interpretation of these observations is that the generation of mature snoRNA present within the parent polyA mRNA through splicing becomes more efficient in response to stress, thereby enhancing their function in modifications of rRNA and snRNA, which ultimately could contribute to selective translation of oxidative stress specific mRNAs. An alternative possibility is that snoRNAs are rapidly degraded under stress conditions, thereby altering their steady-state levels without affecting levels of the spliced parent transcripts.

### Persistent transcription of chaperones after acute stress

Metabolic labeling of RNA allows one to discriminate between alterations in dynamics of RNA production or degradation (Rabani et al., 2011). Conventional RNA-seq does not always reflect transcriptional changes because changed levels of steady-state mRNA can also arise from altered RNA turnover (Bansal et al., 2020; Blatt et al., 2020). To determine the origin of altered transcript levels during stress and recovery as indicated by RNA seq analysis, we *in vivo* labeled nascent mRNAs using 5-ethyl uridine (5-EU) and determined whether there was clear evidence for new transcription of “upregulated” mRNAs using Click-iT, a technique that has been used to distinguish mRNA turnover and *de novo* transcription in several organisms (Battich et al., 2020; Chen et al., 2018; Jao and Salic, 2008; Szabo et al., 2020). For control and acutely stressed cells, we added 5-EU in normal or arsenite-containing medium and collected cells after 3h. To analyze transcription after the stressor had been removed (during recovery), we added 5-EU after 3h of stress and then harvested the cells for RNA isolation (Fig. 4A). We isolated total RNA from all the samples and used the Click-iT Nascent RNA Capture kit to selectively pull down labeled nascent RNA on beads for cDNA synthesis and RNA-Seq. This method captured new transcripts without the need for them to be polyadenylated.

**Figure 4.**
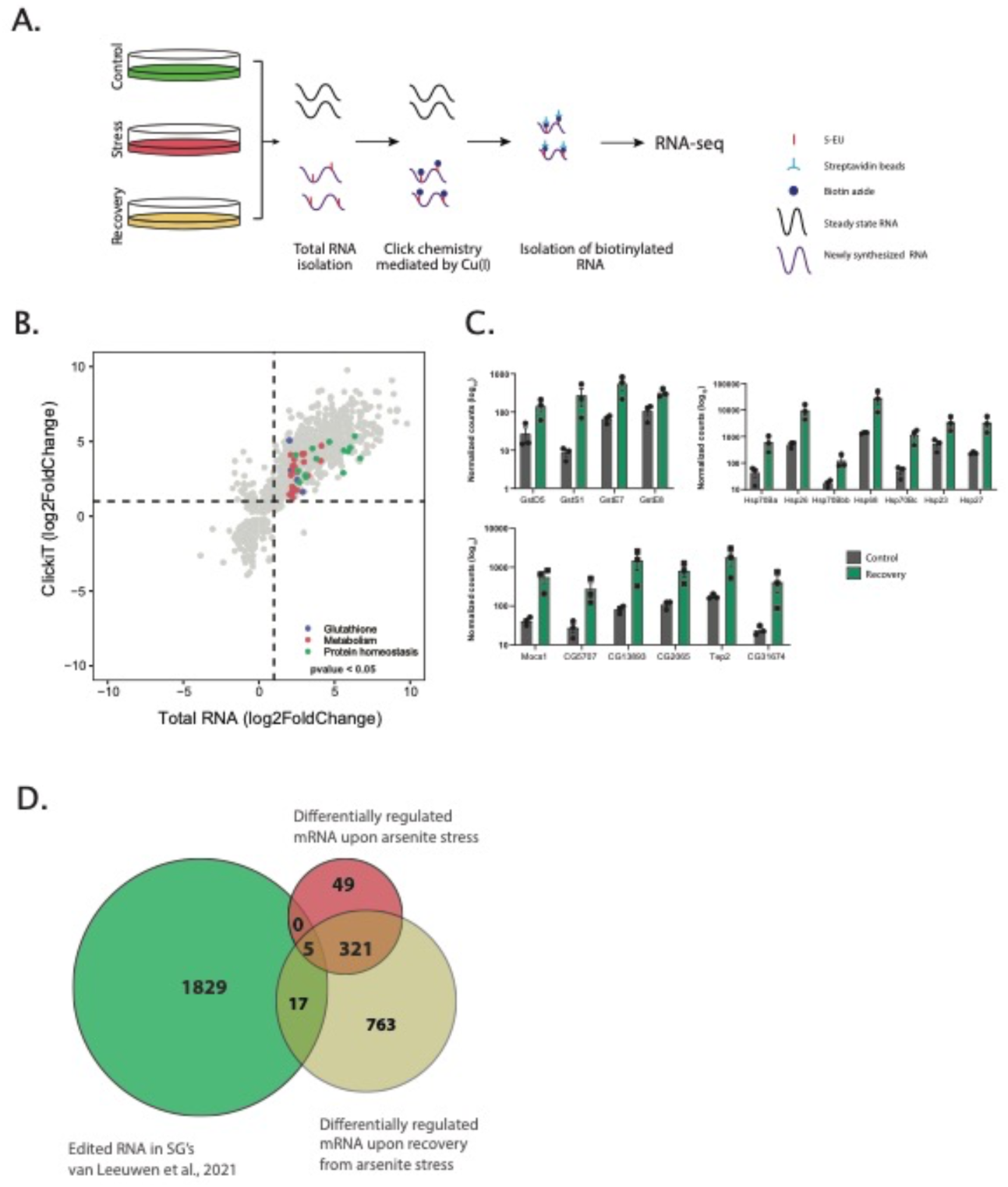
Recovery is characterized by *de novo* transcription. **A.** Schematic for labeling mRNAs using 5-EU. **B.** Enhanced levels of *de novo* synthesized transcripts corresponding to chaperones, GSTs, and genes involved in metabolism during recovery from stress. **C**. Normalized counts of mRNAs during recovery. Transcripts coding for GSTs, chaperones, and metabolism-related genes are shown. **D.** Venn diagrams comparing SG transcriptome (Van Leeuwen et al., 2021) with mRNAs differentially regulated in both stress and recovery.

RNA-seq analysis of the Click-iT captured mRNAs confirmed increased stress-induced transcription of mRNAs whose levels were elevated after stress. Transcripts coding for Hsps, for example, stv, Hsp23, Hsp26, Hsp27, DnaJ-1, Hsp70Bc, Hsp68, Hsp70Ba, etc., were seen as transcriptionally upregulated during recovery (Fig. 4B, C). This observation confirms that new transcription of chaperones occurs during acute stress and continues for a substantial period during recovery from stress. Interestingly, almost all the transcripts which were upregulated in stress and recovery are predicted to be excluded from the SGs (Fig. 4D). Out of the 1856 transcripts reported as present in SGs in *Drosophila* stress granules (Van Leeuwen et al., 2021), we found that only 5 transcripts were included among the stress-regulated mRNAs that we identified (Supplementary File 3). Similarly, only 22, corresponding to 1.2% differentially regulated transcripts in recovery, were found to be present in the reported collection of SG-associated mRNAs (Fig. 4D). This comparison reveals that the mRNAs that are upregulated during stress are excluded from SGs, suggesting that they either have roles in translational repression or encode factors that are translated during stress and recovery.

### Oxidative stress transcriptional response is uncoupled from SG assembly

Given that stress induces both stress-granules and new transcription, we were interested to know whether the assembly of SGs contributed to signaling transcription of at least a significant subset of target mRNAs, as has been proposed following viral infection (Alam and Kennedy, 2019; McCormick and Khaperskyy, 2017; Tsai and Lloyd, 2014). To address this outstanding question, we asked how disrupting SG assembly would affect stress-induced transcription.

The SG protein Rin/G3BP is a primary nucleator of SGs, whose knockdown prevents SG assembly in response to starvation in *Drosophila* S2 cells (Aguilera-Gomez et al., 2017) as well as during several other conditions in different mammalian cell lines (Lee et al., 2020; Sanders et al., 2020; Yang et al., 2020). Apart from its role in SG assembly, the housekeeping functions of Rin/G3BP involve binding to RNA and regulating selective protein synthesis during oxidative stress via mRNA partitioning (Laver et al., 2020; Somasekharan et al., 2020). We used dsRNA-mediated RNAi to knock down the levels of *Rin* in S2 cells and independently assessed the effect of this perturbation on SG granule assembly as well as on stress-induced transcription.

Experimental cells treated with dsRNA targeting endogenous Rin mRNA showed reduced levels of Rin protein compared to mock control cells (treated with dsRNA targeting GFP) (Fig. 5A). In mock control cells, arsenite exposure robustly induced Atx2- and Rin- containing SGs (Fig. 5B). In contrast, and as predicted, *Rin*-RNAi treated cells with reduced *Rin* mRNA and protein (Fig 5A) were unable to form SGs (Fig. 5B). To test whether the inability to form SGs affected the transcriptional response to stress, we used RNA-seq to determine and analyse transcriptomes in control and *Rin* RNAi cells exposed to arsenite as described previously (Fig 2A). Transcriptomes for three control and three *Rin* RNAi replicates showed high internal correlation coefficients within each group (between 0.992 and 1.0), demonstrating high similarity among biological replicates within each condition. However, and remarkably, similar levels of correlation were also seen across groups: indeed, mock and *Rin* RNAi transcriptomes were largely indistinguishable (Supplementary Fig. 4A). This observation suggests that *Rin* knockdown, which prevents normal SG formation, has no significant effect on stress-induced transcription. Consistent with this: (a) transcriptomes of *Rin*-deficient cells following 3 hours of arsenite exposure matched most closely with those of control cells after 3 hours of acute stress (Figure 5 C-ii), and (b) transcriptomes of *Rin*-deficient cells 3 hours post-stress recovery matched most closely with those of similarly treated controls (Figure 5 C-iii).

**Fig. 5.**
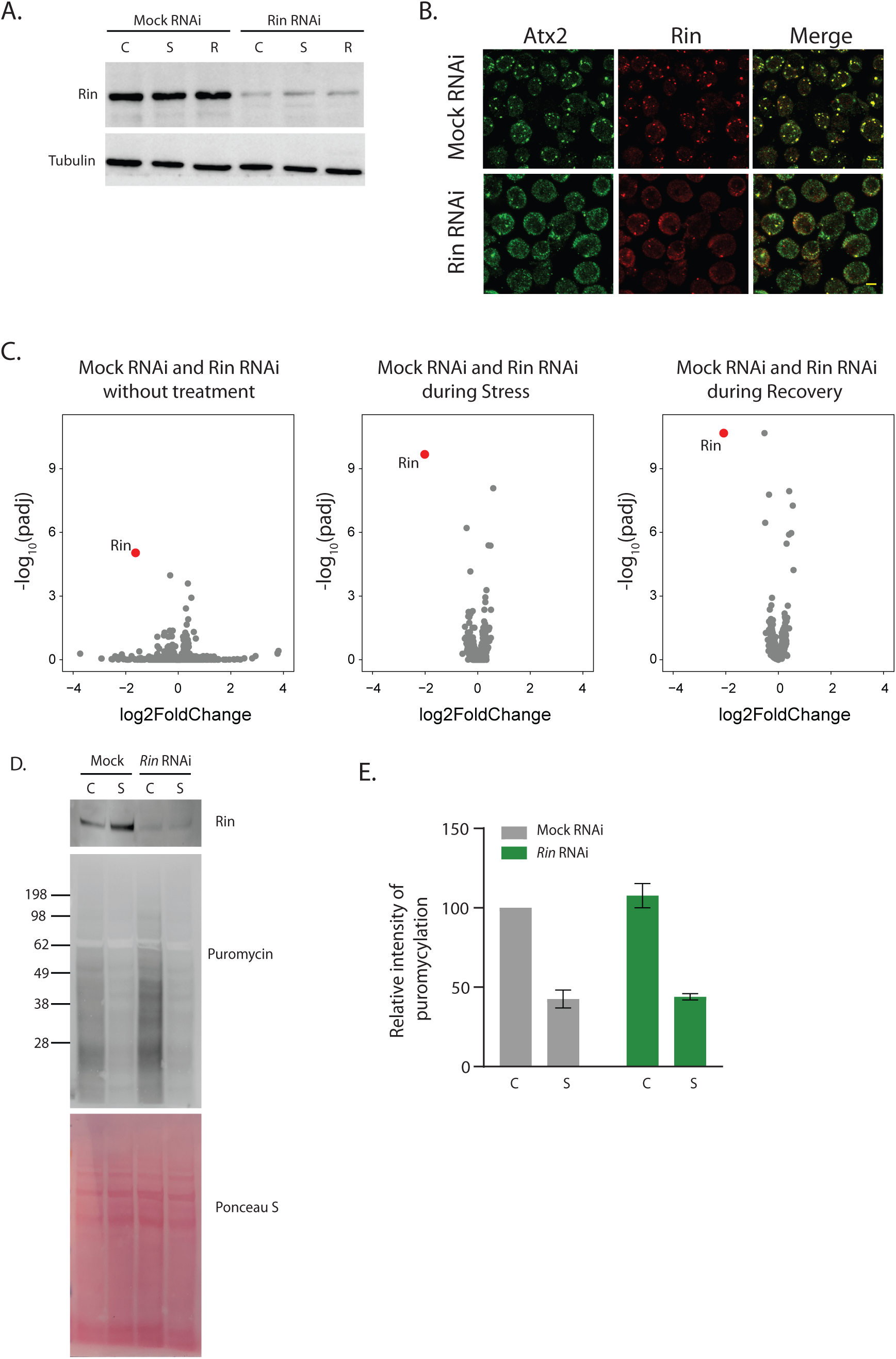
*Rin* knockdown prevents stress granule assembly without altering stress-induced transcription. **A.** Western blot analyses using total cell lysates from mock RNAi and *Rin* RNAi cells under control, stress, and recovery show drastically reduced Rin protein levels in *Rin* RNAi cells. Western blot analyses for tubulin using the same lysates were used as the loading control. Anti-Rin and anti-tubulin antibodies were used. **B.** Arsenite does not cause stress-granule induction in Rin/G3BP deficient cells (after Aguilera-Gomez et al., 2017). Upon stress, both Atx2 and Rin co-localize in granules in mock RNAi cells (i-iii). However, in *Rin* RNAi cells, no granules are assembled upon stress, and Atx2 is diffusely distributed (i’-iii’). Scale bars represent 5 μm. Anti-Atx2 (1:500) and anti-Rin (1:500) antibodies were used for immunofluorescence. **C.** Volcano plots showing the similarity in transcriptomes across mock RNAi and *Rin* RNAi samples with log2FoldChange =1.5 and padj< 0.05. The red dot indicates the levels of Rin in *Rin* RNAi samples. **D.** O-propargyl-puromycin incorporation assays in mock and *Rin* RNAi cells. Western analyses using total cell lysates from mock RNAi and *Rin* RNAi cells under control and stress conditions were used for puromycin incorporation. Anti-puromycin and Anti-Rin antibodies were used at 1:1000 and 1:500 dilutions respectively. A representative blot of four independent experiments is shown. **E.** Bar graphs showing the relative intensity of puromycylation in mock RNAi and *Rin* RNAi cells under control and stress conditions.

Further, volcano plots comparing mock and *Rin* RNAi transcriptomes showed that the transcript levels for all genes remained mostly unchanged following *Rin* knockdown, with the notable exception of *Rin* itself, which was reduced almost four-fold compared to the levels in mock RNAi (Fig. 5C; Supplementary Fig. 4B). The selective effect on *Rin* also confirmed that the effect of *Rin* RNAi was target-specific, with no significant off-target effects. A comparison of the top 100 differentially regulated genes showed no difference among the mock and *Rin* RNAi cells (Supplementary Fig. 4C, Supplementary File 5). These observations suggest that oxidative stress-induced SGs do not have a role in oxidative stress-induced transcription, and they appear to be independent but parallel pathways.

It was notable that neither stress nor *Rin* knockdown had any significant effect on the expression of mRNAs encoding known stress-granule or stress-granule-associated RNA-binding proteins within the time scale of our experiments. Thus, there were no significant changes in the transcript levels for Atx2, Caprin, Cabeza (Fus), Fmr1 (FMRP), Me31B, Pontin (RuvBL1), Reptin (RuvBL2), Ref(2)p (p62/SQSTM1), Rox8 (TIA1), TBPH (TDP43) and Lingerer (UBAP2L) (Supplementary Fig. 4B).

We also used O-propargyl-puromycin incorporation assays to examine whether global translational repression induced by stress was affected under conditions where Rin levels are low, and SGs are not observed. Strikingly, under conditions of reduced levels of Rin where SGs do not form, global translation is still inhibited by stress (Fig. 5D, 5E). This is consistent with previous work in mammalian cells, suggesting that although the translation is widely repressed during stress, only about 5% of mRNA is sequestered within SGs (Khong et al., 2017). Interestingly, levels of Rin showed an increase during stress in mock cells treated with puromycin which further points to another layer of complexity (Fig. 5D).

## DISCUSSION

### The transcriptional response to oxidative stress

Given the importance of oxidative-stress for physiology and disease, there have been relatively few studies of oxidative stress-induced transcription in metazoa (Brown et al., 2014; Zou et al., 2000). However, extensive work in bacteria, plants, and yeast, as well as some in metazoan animal species, have provided important insights (Blevins et al., 2019; He et al., 2018; Reichmann et al., 2018; Wohlbach et al., 2009). Global transcriptional changes that occur during recovery following stress remain even relatively sparsely studied (Sørensen et al., 2005). First, that different types of stress can induce overlapping groups of genes, pointing to the principle of cross-tolerance, wherein proteins induced by and that confer protection to heat stress, for instance, may also be similarly regulated and perhaps protective during oxidative stress (Chowdhary et al., 2019; Dahl et al., 2015; Jacobson et al., 2012; Morimoto, 1998). This could, in part, be explained by overlapping cellular effects of stressors: both heat and oxidative stress alter protein folding, and chaperone systems that prevent protein aggregation or promote refolding may be required in both conditions. Moreover, stress-responsive genes could also encode conserved proteins involved in constitutive cellular maintenance (Kültz, 2003; Rebeaud et al., 2020).

#### Specific suites of genes (and functions) induced by and required under oxidative stress

The induction of oxidative stress by arsenite generates reactive oxygen species (ROS) in the cell, which regulates many stress-regulators, including heat shock proteins (Ruiz-Ramos et al., 2009). *Hsp* mRNAs are, in general known to be upregulated during different stress conditions, aging as well as during development (Brown et al., 2014; Colinet et al., 2010; Colinet and Hoffmann, 2010; Michaud et al., 1997; Ruiz-Ramos et al., 2009; Vos et al., 2016; Zou et al., 2000). In *Drosophila*, *Hsps* are also induced during recovery from cold stress (Colinet et al., 2010; Štětina et al., 2015), however, the type of HSPs and the amount of HSPs induced depends on the type of stress (Morano et al., 2012; Zhao et al., 2015). We find a similar upregulation of *Hsp* mRNAs during stress (Supplementary Fig. 3A) as well as during recovery (Fig. 3). Intriguingly, the upregulation of mRNAs during recovery was via active transcription as seen by metabolic labeling (Fig. 4B) and not because of the enhanced stability of mRNAs during recovery. This is a significant finding as it implies the *Hsp* coding mRNAs, which are upregulated during stress may have separate functions than those upregulated during recovery, just like it has been shown for HSP70 in thermotolerant cells (Tian et al., 2021).

Akin to chaperones, GSTs also have a cytoprotective function; for example, they can protect against oxidative damage to DNA and prevent mutations (Allocati et al., 2018; Veal et al., 2002). As seen for *Hsp* mRNAs, we find that several GST mRNAs are upregulated both during stress (Supplementary Fig. 3A) and are actively transcribed during recovery (Figs. 3A, 4B). In yeast, it is known that GSTs are required for cellular resistance to oxidative stress (Veal et al., 2002). There is also the interesting possibility of GSTs being regulators of stress kinases and thus, modulating signal transduction (Adler et al., 1999; Laborde, 2010). Upregulation of transcripts encoding for proteins involved in such cytoprotective functions points to the fact that to attain homeostasis during recovery, a cell needs to prevent protein aggregation (by the action of sHSPs), fold/refold, misfolded proteins (by a harmonious action of HSP70s and HSP40s) and get rid of free radicals generated due to oxidative stress (by synthesizing more GSTs).

Similar to the coding genes, aberrant upregulation of several ncRNAs is observed during stress and disease conditions (Brown et al., 2014; Connerty et al., 2020; Torrent et al., 2018). We find that several ncRNAs are upregulated during stress and recovery, prominent ones being Hsr omega and RNaseMRP:RNA (Fig. 4). During stress, rRNA processing might be affected, and upregulation of RNaseMRP RNA might be a counteractive response during stress. ncRNAs can likely regulate the stability of mRNAs as they bind to several different proteins and modulate their activity by sequestering them away from their sites of action (Lakhotia, 2012). LncRNAs can also act as a sponge for microRNAs, preventing the cleavage of mRNAs whose translation is required during stress and recovery, as well as regulate translation because of complementarity (Lee and Rio, 2015). Although mature Rpl13a mRNA levels are not affected, oxidative stress reportedly leads to upregulation of intronic C/D box snoRNAs present in the *Rpl13a* gene that is required for propagation of oxidative stress whilst their loss affected mitochondrial metabolism and lowered ROS (Lee et al., 2016; Ly et al., 2017; Michel et al., 2011). Akin to the above observation, we also found several snoRNA transcripts significantly reduced during stress and recovery. These studies imply that the differentially regulated snoRNAs might be crucial for oxidative stress response in *Drosophila* cells as well. Several other snoRNAs are also involved in alternative splicing of mRNAs (Bratkovič et al., 2020; Falaleeva et al., 2016; Kishore et al., 2010; Kishore and Stamm, 2006). It is obvious to speculate that the levels of a set of snoRNAs might be regulated via their splicing while another set of snoRNAs might be involved in promoting alternative splicing of mRNAs.

#### General and specific features of the oxidative stress transcriptome in flies

In the current study, we provide an overview of the global transcriptional changes in *Drosophila* S2 cells upon exposure to sodium arsenite stress and subsequent recovery. The results reveal a general increase in the transcription of *Hsp* genes during both stress and recovery, accompanied by increased transcription of genes coding for detoxifying enzymes and several ncRNAs. We also show that knockdown of *Rin* prevents the assembly of SGs during stress and that oxidative stress-induced transcriptional alterations are a completely independent but a parallel event with respect to SG assembly. The number of transcripts that are differentially regulated during recovery is almost three times more than that in stress and belong to several different classes of proteins as compared to the stress, where the transcripts mainly belong to genes coding for proteins involved in stress response or proteolysis. The upregulation of several transcripts involved in the development and metabolic processes during recovery similarly underlines the efforts being made by the cell to restore homeostasis.

#### Significance of analysis of acute stress and recovery transcriptomes and potential functions for specific classes of genes identified

The transcriptional upregulation of various types of chaperones (Fig. 3A, Supplementary Fig. 3A) suggests that apart from their protein folding role, these proteins are also crucial for preventing promiscuous interactions among aggregation-prone proteins by promoting the formation of SGs (Gitter et al., 2013; Gong and Golic, 2006; Štětina et al., 2015). The chaperones could modulate SG formation and disassembly (Alberti et al., 2017; Ganassi et al., 2016; Mateju et al., 2017). HSP70 has also been found to be present in the cores of ring-shaped TDP43 annuli in neurons (Yu et al., 2021). HSP27 also prevents the entrance of FUS into SGs, suggesting that HSP27 may be necessary for the stabilization of the dynamic phase of SGs (Liu et al., 2020). HSP67BC, another small HSP, has been implicated in preventing toxic protein aggregates in *Drosophila* in a HSP70 independent manner (Vos et al., 2016). Similarly, the yeast HSP40s, Ydj1, and Sis1 are important for the disassembly of SGs (Walters et al., 2015). Upregulation of specific chaperone mRNAs during recovery (Fig. 3A) and the concomitant dissolution of arsenite-induced SGs can be likened to the clearance of protein aggregates achieved by overexpression of specific HSPs (Chan et al., 2000; Huen and Chan, 2005; Vendredy et al., 2020; Vos et al., 2016; Warrick et al., 1999; Webster et al., 2019). In fact, pharmacological activation of HSP70 has been shown to ameliorate neurotoxicity caused by aiding the clearance of polyglutamine aggregates (Wang et al., 2013). Upregulation of transcripts of both ATP-dependent and -independent HSP mRNAs (Fig. 4B, C) might implicate a cellular strategy wherein a cell can employ these proteins in clearing aggregates distinctly and more efficiently (Fare and Shorter, 2021); some of these genes may also be involved in long-term stress adaptation (Bijlsma and Loeschcke, 2005; De Bruijn, 2016).

### SG assembly contributes minimally to the transcription of oxidative stress-induced genes

If SG formation is essential for the cellular stress response, blocking its formation should affect the cellular stress response (Lee et al., 2020). Apart from their role in blocking cellular translation, SGs are also known to stimulate transcription of interferons in response to viral infections suggesting that SGs may modulate transcription indirectly (McCormick and Khaperskyy, 2017; Tsai and Lloyd, 2014). We also find that lowering the levels of Rin, thereby preventing SG formation, had no effect on the inhibition of global translation in S2 cells during stress (Fig. 5D). In yeast cells that were deficient in forming SG in response to heat stress, enormously high levels of mRNAs coding for the HSPs (*HSP12* and *HSP104*) and significantly lower levels of genes involved in rRNA processing, part of the RiBi regulon (*PWP1, UTP13,* and *DIP2*) were found (Yang et al., 2014). The authors opined that this increase or decrease in specific mRNAs levels could be due to alteration in transcription kinetics or altered mRNA stability. In contrast, we find that lowering Rin levels hence inhibiting the formation of SGs, does not affect transcription under stress (Fig. 5). This is surprising because Rin has several housekeeping functions apart from being essential for SG condensation, but it may also not be required in specific cells (Baumgartner et al., 2013; Buddika et al., 2020; Guillén-Boixet et al., 2020; Kedersha et al., 2016; Laver et al., 2020; Pazman et al., 2000; Sanders et al., 2020; Yang et al., 2020). Strikingly, comparative transcriptome analysis between mock and *Rin* RNAi cells revealed no change in the type of differentially regulated transcripts nor any significant alterations of fold changes in expression of individual mRNAs during stress and recovery (Fig. 5). The differentially regulated transcripts in stress and recovery are also excluded from SGs (Fig. 4D), which also implies that the arsenite-induced SG assembly and transcriptional alterations are parallel but independent events. There might be several underlying layers of cellular intricacies that might link these two events.

Transcription of stress-responsive genes serves a crucial role in implementing a rapid and robust stress response (Vihervaara et al., 2018, 2017). Upon stress removal, the SGs dissolve, and cap-dependent translation begins as suggested by the loss of eIF2a phosphorylation; however, if these recovery responses are attributed to the reversal of transcriptional changes that had occurred during stress is not known (Fig. 6). Further, if transcriptional dysregulation during recovery plays any role in SG dissolution remains unknown. Ultimately, comparing stress and recovery responses as a continuum but not in isolation is crucial in dissecting these two phenomena with exact opposite consequences to cellular homeostasis.

**Fig. 6:**
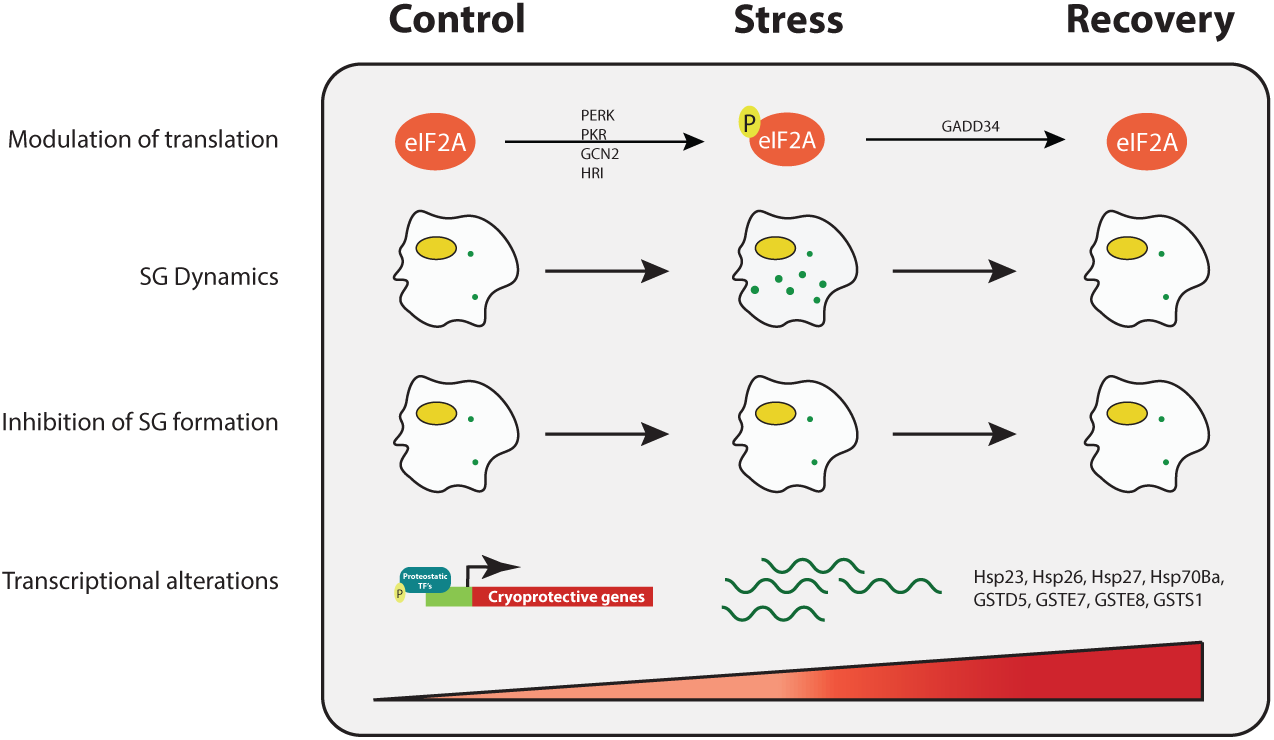
A model depicting the various cellular changes taking place during stress and subsequent recovery. eIF2α gets phosphorylated at serine 51 during stress by the action of any of the four kinases, leading to a block in cap-dependent translation. Upon the removal of stress, phosphorylation is lost, and cap-dependent translation is restored. This is concomitant with assembly and clearance of SGs respectively during stress and recovery, as well as with increased transcription of cytoprotective genes such as HSPs and GSTs mediated by proteostatic transcription factors (HSF1, FOXO, NRF2, etc). However, when the formation of SG is prevented by lowering down levels of Rin, there is no change in the transcription of the cytoprotective genes.

We propose that alterations in oxidative transcriptional response are a cellular response against long-term chronic stress. At the same time, the assembly of SGs is an immediate effect to counter stress (Fig. 6). However, it remains to be elucidated what the involvement is of the genes that are differentially regulated during recovery upon the dissolution of SGs? Since *Rin* RNAi cells do not form visible SGs, it raises several important questions, (i) although mRNA is devoid of ribosomes, what is their fate? (ii) what is the status of the global proteome when SG assembly has been prevented? Further studies need to be undertaken to address these questions.

## ACKNOWLEDGMENTS

We thank Prof. K. VijayRaghavan for his valuable feedback and suggestions during the course of this work. We also thank the members of the Ramaswami, and Bakthavachalu labs, as well as Prof. Roy Parker (HHMI and University of Colorado Boulder) for useful discussions and/or comments on the manuscript. We thank Prof. Elizabeth Gavis (Princeton University, USA) and Dr. Nicholas Sokol (Indiana University, USA) for sharing Rasputin and Rox8 antibodies with us. We acknowledge Drosophila Genomics Resource Centre (supported by NIH grant 2P40OC010949) for *Drosophila* S2 cells. The work was supported by a Science Foundation Ireland (SFI) Investigator grant to MR, and an Irish Council Postgraduate Fellowship to GT. The work is partly supported by the DBT/Wellcome Trust India Alliance Fellowship (IA/I/19/1/504286) awarded to BB, who is also a recipient of the SERB-STAR award (STR/2020/000056). We acknowledge the support from INSA Young Scientist Project (INSA/SP/YSP/143/2017) (AS), from SERB to MR from a collaborative VAJRA award to Dr. Raghu Padinjat, and a CSIR fellowship (DJ). We thank Dr. Awadhesh Pandit from the NCBS NGS facility for help with RNA-seq.

## Methods

### Cell culture and treatments

*Drosophila* S2R+ cells were obtained from DGRC and cultured in the semi-adhering state in Schneider’s medium (S2 medium) with 10% FBS, penicillin, and streptomycin at 25^0^C. Cells were maintained at 50% confluency in fresh S2 media for atleast 24h before being used for stress experiments.

To induce stress, cells were subjected to 0.5mM sodium arsenite in S2 media for 3 hours at room temperature on a rocking shaker. After 3 hours, arsenite containing S2 media was removed by centrifugation at 2000 rpm for 5’, and the cells were washed three times with fresh media and kept for recovery in fresh complete S2 media. Recovering cells were kept on the rocking shaker for an additional 3h at room temperature.

For labeling with 5-EU, 200um EU was added as indicated in Fig 4. Briefly, for labeling transcripts under control and stress conditions, 5-EU either in a normal medium or in a medium containing 0.5mM sodium arsenite at the start of the 3h stress regime. Cells were then washed and harvested for RNA isolation. For labeling transcripts during recovery, cells were initially stressed with 0.5mM sodium arsenite for 3h and then were washed three times with fresh S2 media. 5-EU was added at the start of the 3h recovery period, after which cells were harvested for RNA isolation.

### RNA isolation and RNA Seq

After stress and recovery, RNA was isolated using TRIzol reagent (Invitrogen, USA) as per the manufacturer’s protocol. RNA concentration was measured using a Qubit RNA assay kit in Qubit 2.0 Fluorometer (Life Technologies, USA). RNA integrity was confirmed using the RNA Nano 6000 assay kit of the Bioanalyzer 2100 system (Agilent Technologies, USA). Poly(A) enriched mRNA library was made using TruSeq RNA Library Preparation Kit V2 (RS-122-2001) and sequenced using HiSeq SR Rapid Cluster Kit v2 (GD-402-4002) to generate 1X50 single-end reads on Illumina HiSeq2500 sequencing platform.

### *In silico* analysis

For transcriptome analysis, all sequencing reads obtained post adaptor removal had a mean quality score (Q-Score) >= 37, so no trimming was required. All the further downstream analyses were performed on this high-quality data. For read mapping, the reference genome and gene model annotation files of *D. melanogaster,* version dm6 were downloaded from the UCSC genome browser. Single-end processed reads were aligned to the reference genome using STAR v2.5.3 with default parameters. HTSeq-count v0.11.2 and the “-s reverse” option were used to count the read numbers mapped to each gene before differential gene expression analysis. Differential gene expression among samples was performed using the DESeq2 package. The data is available with the assigned GEO accession # GSE178464. For Gene Ontology analysis of differentially expressed genes under stress and recovery, enriched GO terms with <0.05 *p*-value were identified using BINGO plug-in at Cytoscape (v.3.8.0) and the enriched GO terms were shown in heatmap via MeV (v.4.9.0). For granule counting, Cellprofiler was used. On an average, 140 cells per field were acquired. The number of granules observed in each cell were tabulated for control, stress and recovered cells. Outliers were determined by calculating the number of granules that were 1.5 standard deviations above and below the mean and were thus excluded from further analysis. The values were plotted as average number of granules observed as well as the numbers of granules per cell. On applying Mann-Whitney statistics, it was observed that there was a significant difference in the number of granules between stressed and recovered cells (p < 0.05).

### Immunostaining and fluorescence

Immunostaining was performed as described earlier (Bakthavachalu, Huelsmeier et al., 2018). Briefly, S2R+ cells were grown in T25 flasks to almost 70-80% confluency. Stress and recovery experiments were performed as described above. Cells were fixed with 4% paraformaldehyde for 10 min, followed by permeabilization with 0.05% Triton-X-100 for 10 min. This was followed by blocking with 1% BSA for 30 min. The cells were then incubated with antibodies against Atx2 (1:500), Rox8 (1:1000) and Rin (1:500), followed by probing with 1:1000 dilution of Alexa Fluor 488, 568 and 647 (Abcam) secondary antibodies respectively. Confocal imaging was done using the PALPON 60x/1.42 oil objective of the Olympus FV3000 microscope. Images were processed using ImageJ software.

### Double-stranded (ds) RNA generation

Mock and *Rin* RNAi were performed using dsRNA produced by *in vitro* transcription (IVT). For mock, we utilized GFP open reading frame as the target site. RNAi target sites were chosen using the SnapDragon tool (https://fgr.hms.harvard.edu/snapdragon) (Hu et al., 2017). PCR generated DNA templates containing the T7 promoter sequence at both ends were used as IVT template for dsRNA synthesis using Megascript T7 High Yield Transcription kit (Invitrogen). The primer details are provided in Supplementary File 6.

### dsRNA transfection into cells for RNAi experiments

Wild type *Drosophila* S2 cells were depleted for *Rin* mRNA by double-stranded (ds) RNAi. Briefly, 0.5 million cells were transfected with 5μg of dsRNA. After 48h of the first round of transfections, cells were again transfected with 5μg of dsRNA. After 96 hours of the first transfection, cells were analyzed for the knockdown of Rin, and RNA was isolated using TRIzol reagent (Invitrogen) as per the manufacturer’s protocol. Illumina library was prepared from Poly(A) enriched mRNA using TruSeq RNA Library Preparation Kit V2 (RS-122-2001) and sequenced using HiSeq SR Rapid Cluster Kit v2 (GD-402-4002) to generate 1X50 single-end reads.

For puromycylation assays, after 93 hours of mock and *Rin* dsRNA transfections, stress was induced as previously described above. Puromycin was added during the final 15 minutes at a final concentration of 4μg /mL. Cells were then immediately harvested and analysed for the knockdown of Rin, and for puromycin incorporation.

### Protein isolation and Western analysis

Mock RNAi and *Rin* RNAi cells were maintained, stressed, and recovered as mentioned above. Total protein isolation was performed as described earlier (Sudhakaran et al., 2014). Briefly, 0.2 million cells were pelleted and resuspended in 50ul lysis buffer [25mM Tris HCl (pH7.5), 150mM NaCl, 10% (v/v) glycerol, 1mM EDTA, 1mM DTT, 0.5% Nonidet P-40 and complete protease inhibitor tablets from Roche) and incubated at 4^0^C for 30 min with intermittent vortexing. The lysate was then spun at 4^0^C at 13000 rpm for 30 min. The supernatant was collected, and protein was quantified using Nanodrop. For puromycylation, cells were lysed and normalised for protein concentration using the Bio-Rad Protein Assay Kit (Bio-Rad Laboratories, Inc. # 5000001) and a spectrophotometer. Westerns blots were performed using rabbit anti-Rin (1:1000), rabbit anti-phospho-eIF2α (1:1000, 9721L CST), rabbit anti-eIF2α (1:1000, SAB4500729-100UG), mouse anti-puromycin (1:2000, MABE343 Sigma-Aldrich) and mouse anti-tubulin (1:2000, E7c DSHB). Goat anti-Rabbit HRP (sc-2004) and goat anti-mouse HRP (sc-2005) HRP-conjugated secondary antibodies were used at 1:10000 dilution.

---

**Supplementary Fig. 1:**
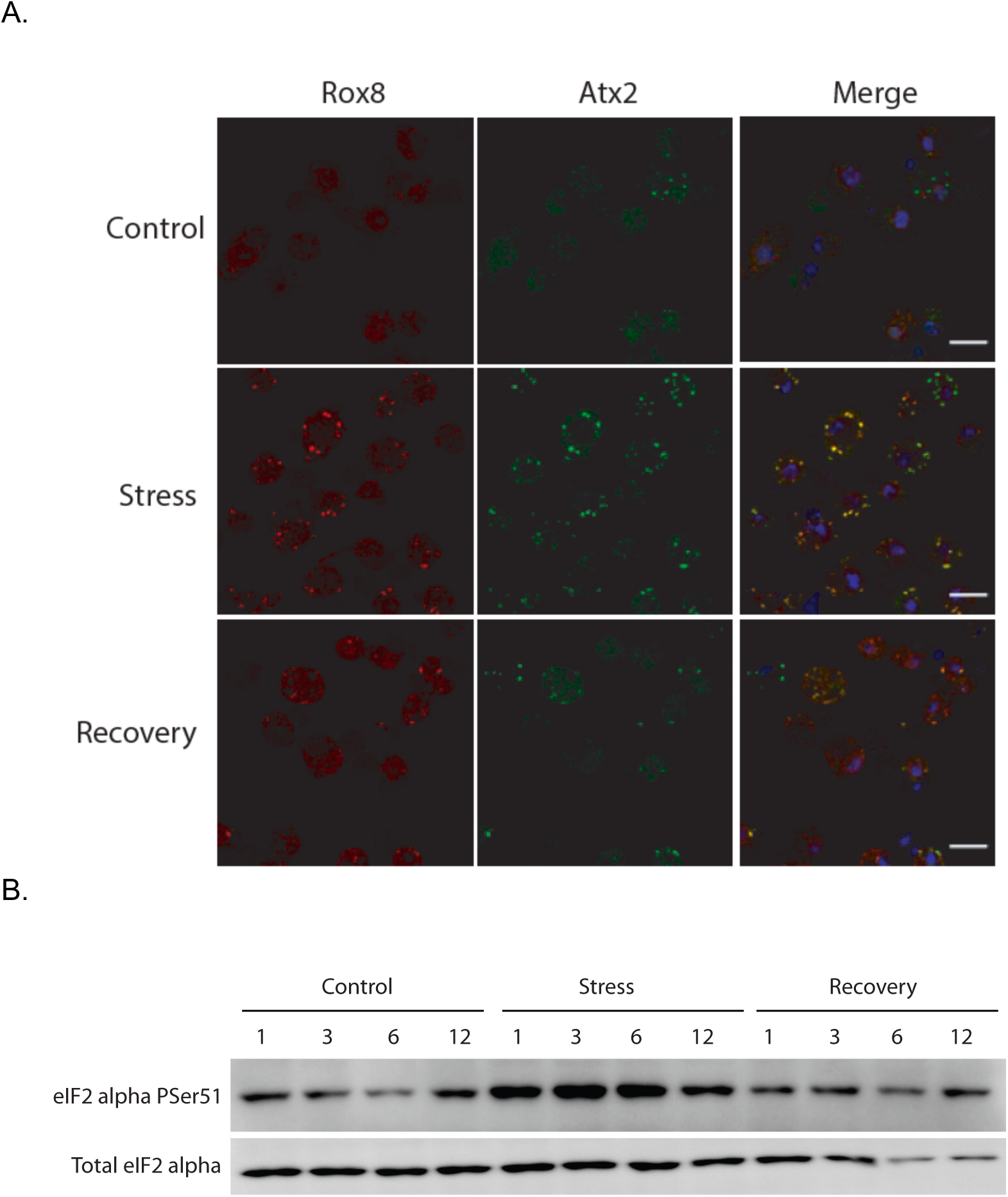
**A.** Control stressed and recovered cells were stained with Hoechst33342 (dilution 1:1000) for staining the nucleus and probed for Atx2 and Rox8 which have been shown to colocalize in granules upon stress (Bakthavachalu et al., 2018; Farny et al., 2009). Images were acquired on Olympus FV3000 6 lasers using a PLAPON60XO 1.42 NA objective at 1024 x 1024 resolution. Two fields of cells were acquired for each condition. Z projections of slices of cells acquired with a 1 μm step size were used for granule quantification. The number of cells and the granules present in the cells were quantified using CellProfiler. The punctae formed by Rox8 were annotated as granules and Hoechst33342 staining was used for identifying cells. The cytoplasmic boundary was defined such that punctae observed beyond the plasma membrane were considered as background. Clumped objects were distinguished based on intensity and thresholding for granule quantification was maintained in the range of 0.2 to 1.0. The scale bars represent 10μ. **B.** Levels of total and phosphorylated eIF2α under control, stress and recovery. Total cell lysate was taken at 1h, 3h, 6h and 12h from control, stress and recovery cells and analyzed for serin 51 phosphorylation of eIF2α (upper panel). Lower panel shows the levels of total eIF2α.

**Supplementary Fig. 2:**
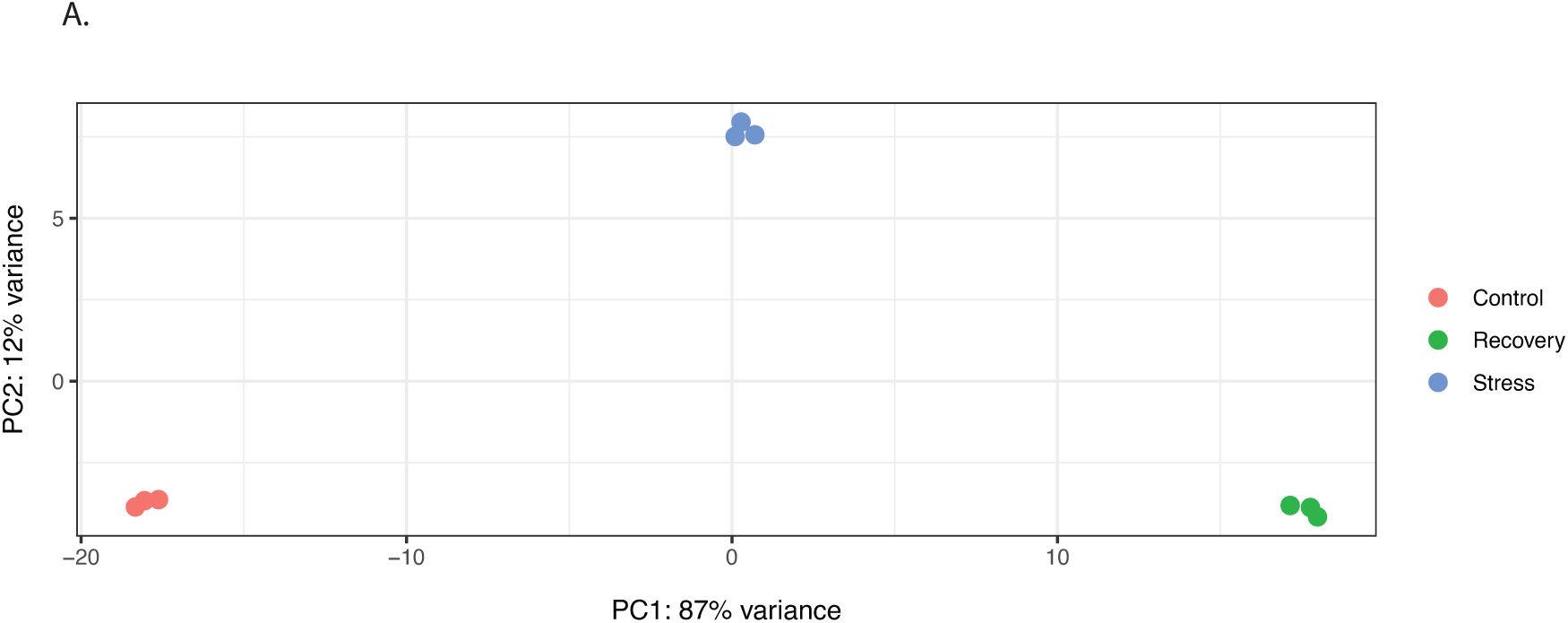
Principal component analysis (PCA) of the data from control, stress and recovery transcriptomes. PCA shows that within the replicate transcriptomes of control, stress and recovery, there is no variation. However, the control, stress and recovery transcriptomes themselves vary a lot among each other.

**Supplemental Fig. 3:**
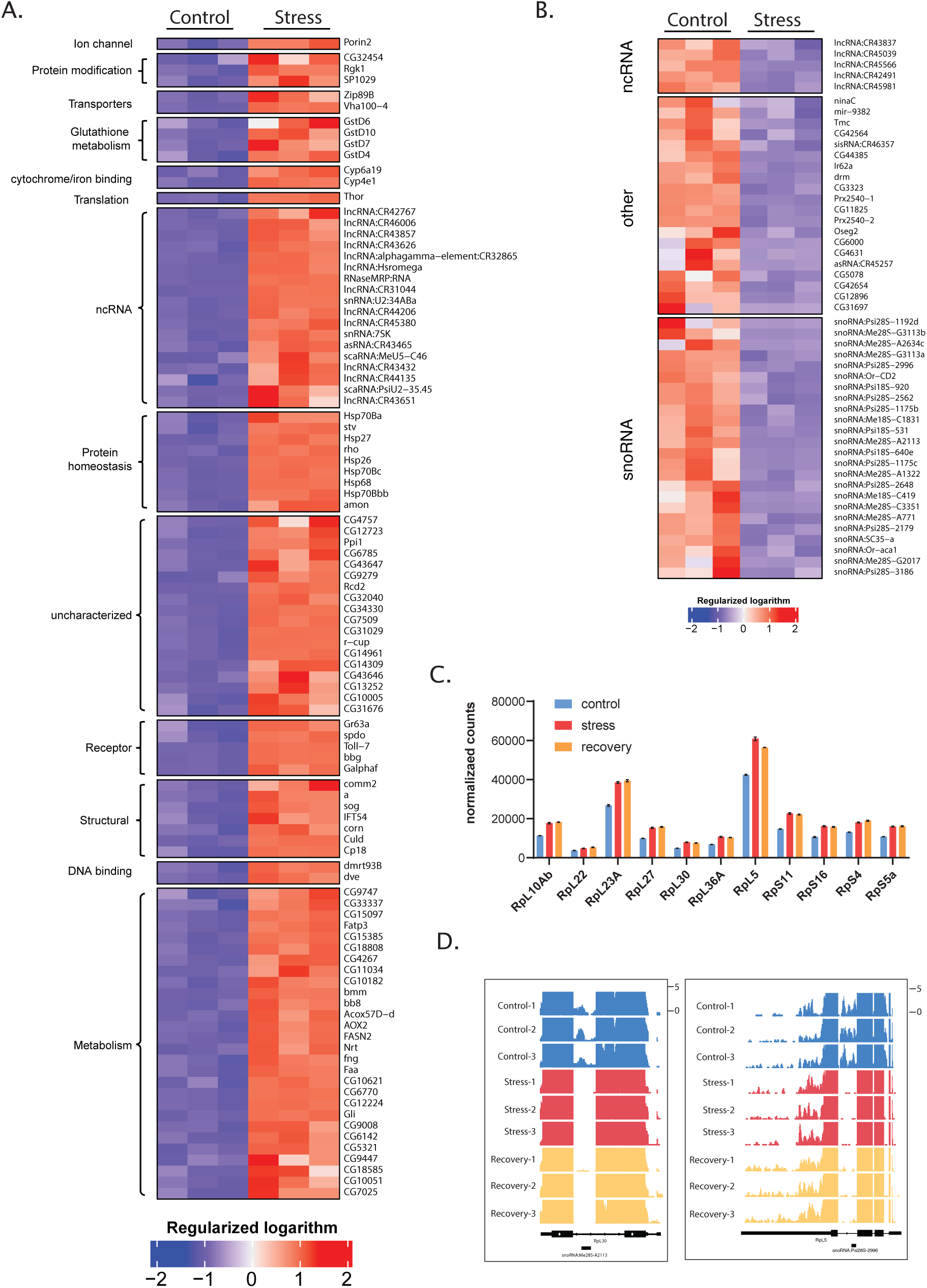
Oxidative stress results in widespread changes in transcription. **A.** Heat map for 100 genes most robustly upregulated following 3-hours of acute stress. Genes are grouped based on predicted cellular functions. The fold induction is indicated in the colour scale below. **B.** Heat map shows a smaller group of mRNAs for which reads are substantially decreased after acute stress. The colour scale bar indicated fold changes represented. **C.** Normalized counts of the parent ribosomal protein genes in control, stress and recovery showing no significant change in the levels. **D.** Bed graphs showing the reads corresponding to snoRNA:Me28S-A2113 (i) and snoRNA:Psi28S-2996 (ii) under control, stress, and recovery. Reads corresponding to the parent genes are also depicted.

**Supplemental Fig. 4:**
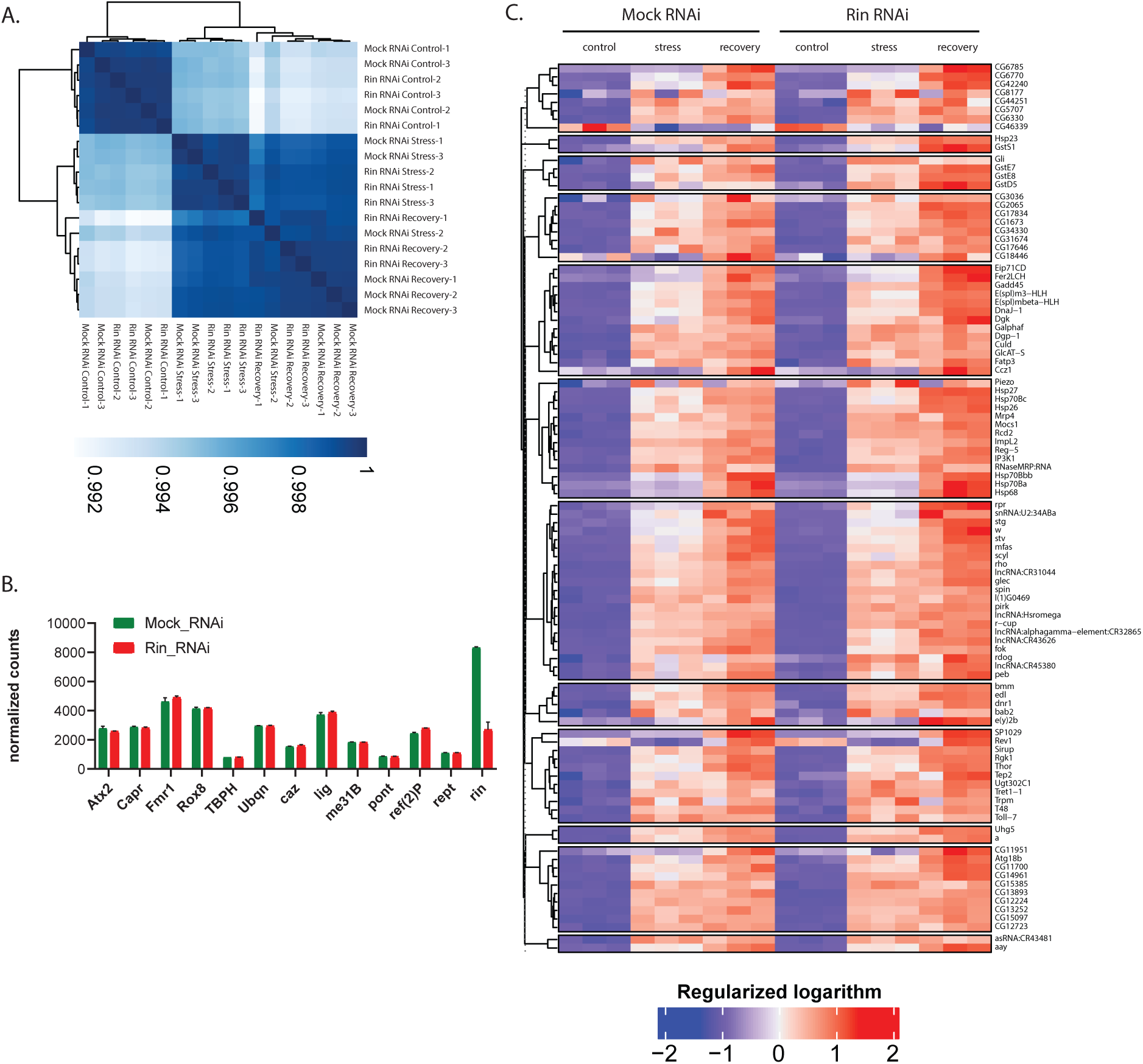
A. Correlation plot depicting the similarities and differences among different transcriptomes. B. Normalized counts of transcripts coding for different SG constituent proteins in mock RNAi and *Rin* RNAi cells. C. Heat maps depicting transcriptional alterations in the top100 genes mock and *Rin* RNAi cells under control, stress, and recovery with a cutoff of log2 fold.

## Notes

### Competing Interest Statement

The authors have declared no competing interest.

